# Whole genome data reveal the complex history of a diverse ecological community

**DOI:** 10.1101/233759

**Authors:** Lynsey Bunnefeld, Jack Hearn, Graham Stone, Konrad Lohse

## Abstract

How widespread ecological communities assemble remains a key question in ecology. Trophic interactions between widespread species may reflect a shared population history, or ecological sorting of local pools of species with very different population histories. Which scenario applies is central to the stability of trophic associations, and the potential for coevolution between species. Here we show how alternative community assembly hypotheses can be discriminated using whole genome data for component species, and provide a novel likelihood framework that overcomes current limitations in formal comparison of multispecies histories. We illustrate our approach by inferring the assembly history of a Western Palaearctic community of insect herbivores and parasitoid natural enemies, trophic groups that together comprise 50% of terrestrial species. We reject models of co-dispersal from a shared origin, and of delayed enemy pursuit of their herbivore hosts, arguing against herbivore attainment of ‘enemy-free space’. The community-wide distribution of species expansion times is also incompatible with a random, neutral model of assembly. Instead, we reveal a complex assembly history of single- and multi-species range expansions through the Pleistocene from different directions and over a range of timescales. Our results suggest substantial turnover in species associations, and argue against tight coevolution in this system. The approach we illustrate is widely applicable to natural communities of non-model species, and makes it possible to reveal the historical backdrop against which selection acts.

---

The mode and timescale over which complex ecological communities assemble remain largely unknown [1]. Do such communities arise once, and spread through co-dispersal of component species, such that ecological interactions between species are sustained though time and space? Or do they arise repeatedly through local sorting of the same pool of widespread species, with little stability of ecological interactions at the population level? Alternatively, community assembly may be an entirely random and ecologically neutral process. The answer is central to understanding how traits associated with species interactions evolve: while co-dispersal allows co-adaptation among interacting species, the same is not expected for community assembly by ecological sorting or neutral processes, in which species can have very discordant population histories.

Concordant population histories are expected in obligate species associations, such as those between plants and specialist pollinators, and these commonly show both coevolution of associated traits and co-diversification of the lineages involved [2, 3]. However, much of terrestrial diversity is found in communities dominated by less specific interactions between guilds of species, for which either co-dispersal or ecological sorting are plausible assembly mechanisms. These are exemplified by the rich insect communities associated with temperate trees, in which widespread herbivores are commonly attacked by a consistent set of parasitoid enemies [4, 5]. Both herbivores and parasitoids in such systems show traits that structure trophic interactions [6, 7, 8], but the extent to which these represent co-adaptations or the results of ecological sorting remains little-understood.

Here we assess the evidence for four alternative models (Fig. 1a-d) in the assembly of an exemplar community of oak-associated insects, comprising herbivorous insect hosts and parasitoid and inquiline natural enemies (Fig. 1, see legend). These models include strict co-dispersal (simultaneous range expansion), host tracking (parasitoid pursuit of their hosts) and ecological sorting (local recruitment of parasitoids from alternative hosts), each of which makes contrasting predictions for spatial patterns of genetic variation within and among species. We also evaluate the support for an alternative null model of assembly under ecological neutrality that assumes that species are trophically equivalent [9, 10].

**Fig 1.**
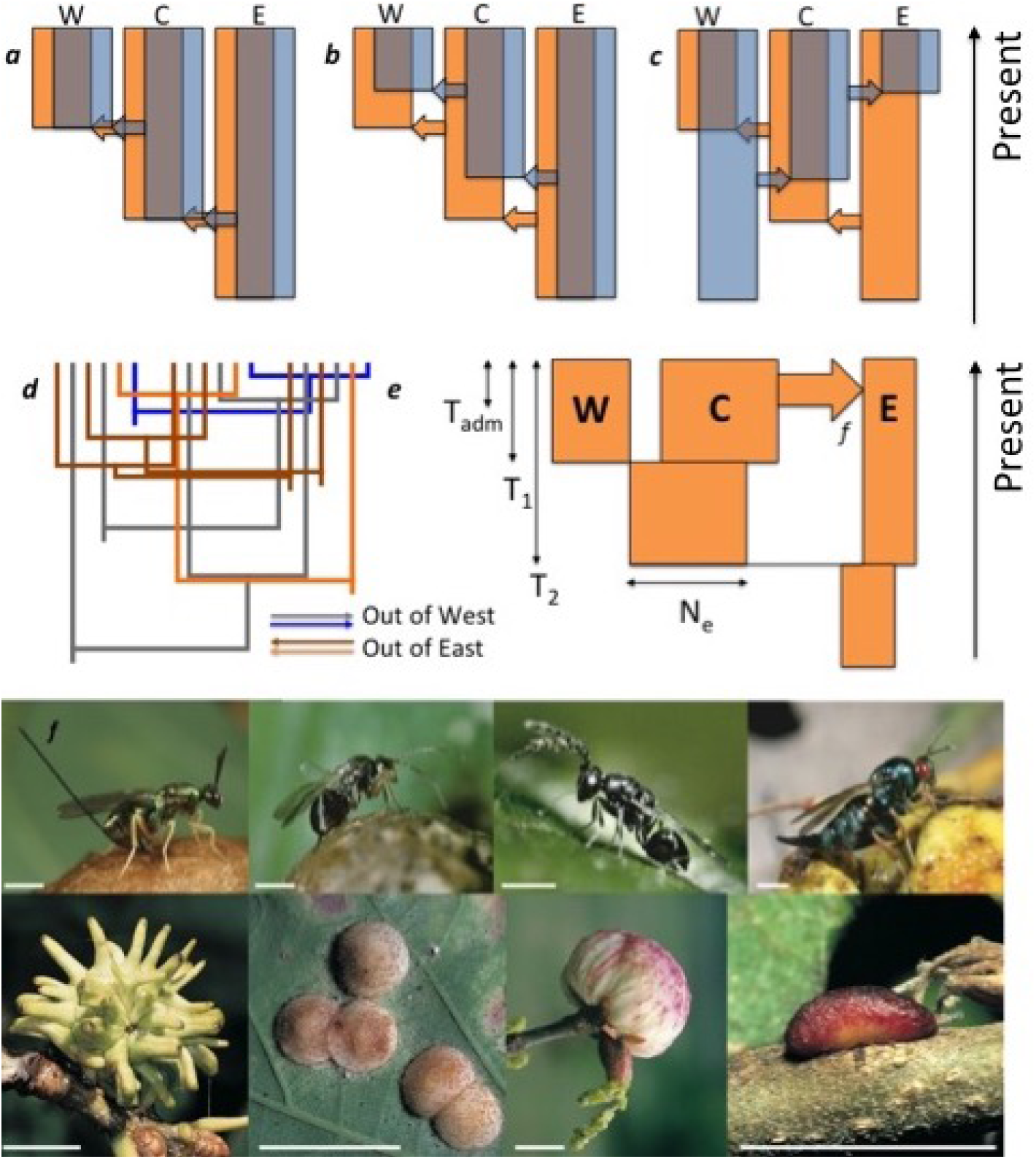
Population histories under alternative hypotheses of community assembly. a) Under strict co-dispersal species disperse between refugia together from a shared origin and their population histories show the same topology and timing of population splits. b) Host tracking predicts that hosts and parasitoids share a common geographic origin, but allows parasitoids to follow after their hosts. Host tracking is significant ecologically because hosts can achieve a measure of enemy-free space [16] which decouples coevolutionary interactions between trophic levels. Enemy escape has been seen over ecological timescales [17] but has rarely been studied in the context of population histories [16]. c) Ecological sorting results in the same set of interacting species in each refuge, but allows highly discordant patterns of range expansion from different origins. d) Under an ecological and biogeographic null model, expansion times are random draws from a single community-wide distribution. The figure shows a six species community with two parasitoids (orange) and one gallwasp host (brown) expanding out of the East and two parasitoids (grey) and one gallwasp (blue) expanding out of the West. Split times (*T*_1_ and *T*_2_) are drawn randomly from an exponential distribution. e) The demographic history of each species is captured by a seven parameter model. We allowed a separate population size (*N_e_*) for each refugial population (3 parameters). In this example, *N_E_* < *N_W_* < *N_C_* (shown by width of bars & where *N_E_* is the *N_e_* for for the Eastern refugium, and so on) and an instantaneous admixture event (shown by a horizontal arrow) at *T_adm_* transfers a fraction *f* of the Central population into the East. f) Exemplar members of the oak gall community. Galls induced by the four oak gallwasp species (bottom) are attacked by a range of parasitoid wasps (top). Natural enemies from left to right *(Synergus umbraculus* is an inquiline oak gallwasp that inhabits galls induced by other oak gallwasps and will be grouped with the other parasitoid species for brevity hereafter): *Torymus auratus, Synergus umbraculus, Eurytoma brunniventris* & *Ormyrus nitidulus.* Galls from left to right: *Andricus grossulariae, Neuroterus quercusbaccarum, Biorhiza pallida*, & *Pseudoneuroterus saliens.* Scale bar is 1 mm on the top row & 1 cm on the bottom row.

Our exemplar community comprises oak cynipid gallwasp herbivores and their chalcid parasitoid wasp natural enemies, which form a set of interacting species whose distributions span the Western Palaearctic from Iberia to Iran with three main refugial centres of intraspecific genetic diversity. Associated parasitoids only attack cynipid galls, making this system ecologically closed and hence suitable for analysis in isolation [4]. Single-species analyses of community members [11, 12, 13, 14] have found westwards declines in genetic diversity that support expansion into Europe from Western Asia, an ‘out of the East’ pattern that is concordant with inferred divergence of Western Palaearctic gallwasp lineages in Western Asia 5-7 million years ago [15]. However, the extent to which this is true of the whole community, and particularly the parasitoids, remains unclear [16]

We test the predictions of each model of community assembly using whole genome sequence (WGS) data for 13 species (four gallwasp herbivores and nine parasitoids) of the oak gall community, each assembled *de novo* and sampled across three Pleistocene refugial populations spanning the Western Palaearctic: Iberia (West), the Balkans (Centre) and Iran (East) (Table S1). For each species and each refugium, we generated WGS data for two males whose haploidy greatly facilitates bioinformatic and population genetic analyses. We infer an explicit history of divergence and admixture between refugia for each species using a computationally efficient composite likelihood framework [18] based on the joint occurrence of mutations in sequence blocks [19]. Analysing WGS data in a blockwise manner overcomes the limitations of previous comparative studies which have either ignored demography altogether [20, 21] or lacked power because of the small number of loci available [16]. We conduct a formal comparison of demographic histories across species and between trophic levels to address the following questions:

1. To what extent do community members share a common origin, direction, and timescale of range expansion across the Western Palaearctic?
2. Did gallwasp herbivores disperse before their parasitoids, and so achieve “enemy-free space”?
3. Is there evidence of dispersal between refugia following their initial colonisation, compatible with multiple waves of range expansion?
4. Can the timing of expansion events across the full set of species be explained by a neutral community assembly model?

We find a diversity of population histories in oak gallwasps and their parasitoids. While most species dispersed into Europe from the East, four species joined the community following dispersal from the West. Evidence of gene flow between refugial populations implies multiple range expansion events and the potential for dynamic community evolution on a continental scale. Species also vary in the timing of their range expansions, with initial divergences between refugial populations dating from 100 thousand years (ky) to 1 million years (my) ago. A likelihood based comparison of these histories allows us to reject both strict co-dispersal of this community and herbivore occupation of enemy free space, two paradigmatic models of deterministic community assembly. Hosts as a guild did not disperse before their parasitoids, providing no evidence for (perfect) enemy escape. However, our results also argue against an ecologically neutral null model of community assembly that views the histories of individual species as random draws from a simple community-wide distribution. Instead, we identify a complex history for this community including a mixture of idiosyncratic range expansions combined with multi-species pulses of range expansion that correspond to climatic shifts during the Pleistocene. The variation of demographic histories we uncover across species suggests an extensive role for ecological sorting. The fact that population histories are largely uncoupled across members of the oak gall community implies that any host-parasitoid coevolution is likely to have been diffuse which is consistent with the observation that this and other temperate communities centered around insect herbivores are dominated by generalist parasitoids.

## Results

We generated whole genome sequence data (100 base paired-end Illumina) for a total of 75 individual haploid male wasps. For each of the 13 species we sampled two individuals from each of the three refugia (West, Centre and East; Table S2). Reads for each species were combined to assemble reference genomes *de novo* (Table S1) and mapped back to generate variation data.

### Defining demographic model space

Our initial aim was to infer the history of longitudinal range expansions into and admixture between refugial populations for each species. We defined a space of plausible demographic models that is both biologically realistic, yet small enough to allow for an exhaustive comparison of all possible population relationships. Cross-species comparisons of the pairwise genetic diversity within (*π*) and divergence between (*d_XY_*) refugia provide several immediate insights into the demographic history of the oak-gall community and were used to guide the selection of appropriate models of population history (Supp. Info 1). These summaries indicated that a minimal description of the longitudinal history of each species should include i) differences in effective population size between refugia, ii) divergence between populations and iii) the possibility of admixture between refugia. These processes can be captured by a seven parameter model (Fig. 1e) that includes the effective population size *N_e_* of each refugium (*N_E_*, *N_C_*, *N_W_*), three time parameters describing the divergence between populations (*T*_1_ and *T*_2_) and an instantaneous, unidirectional admixture event at time *T_adm_* that transfers a fraction *f* from a source population into another refugium. To restrict model complexity to a computationally feasible number of parameters, we only considered admixture that involved the older population either as source or sink (Fig. 1e).

For each species, we conducted a full search of model space (a total of 48 full models), which encompasses three possible orders of population divergence, four combinations of ancestral *N_e_* and four possible unidirectional admixture events and determined the best supported model. We also assessed the support for all simpler scenarios nested within these full models, including divergence without admixture (*f* = 0), polytomous divergence from a single ancestral population (*T*_1_ = *T*_2_), and complete panmixia across the range (*T*_1_ = *T*_2_ =0). To get a sense of the likely timescale of community assembly, we converted time estimates into years using a direct mutation rate estimate for *Drosophila melanogaster* [22] of 3.5 * 10^−9^ mutations per base and generation (Table 1).

**Table 1.**
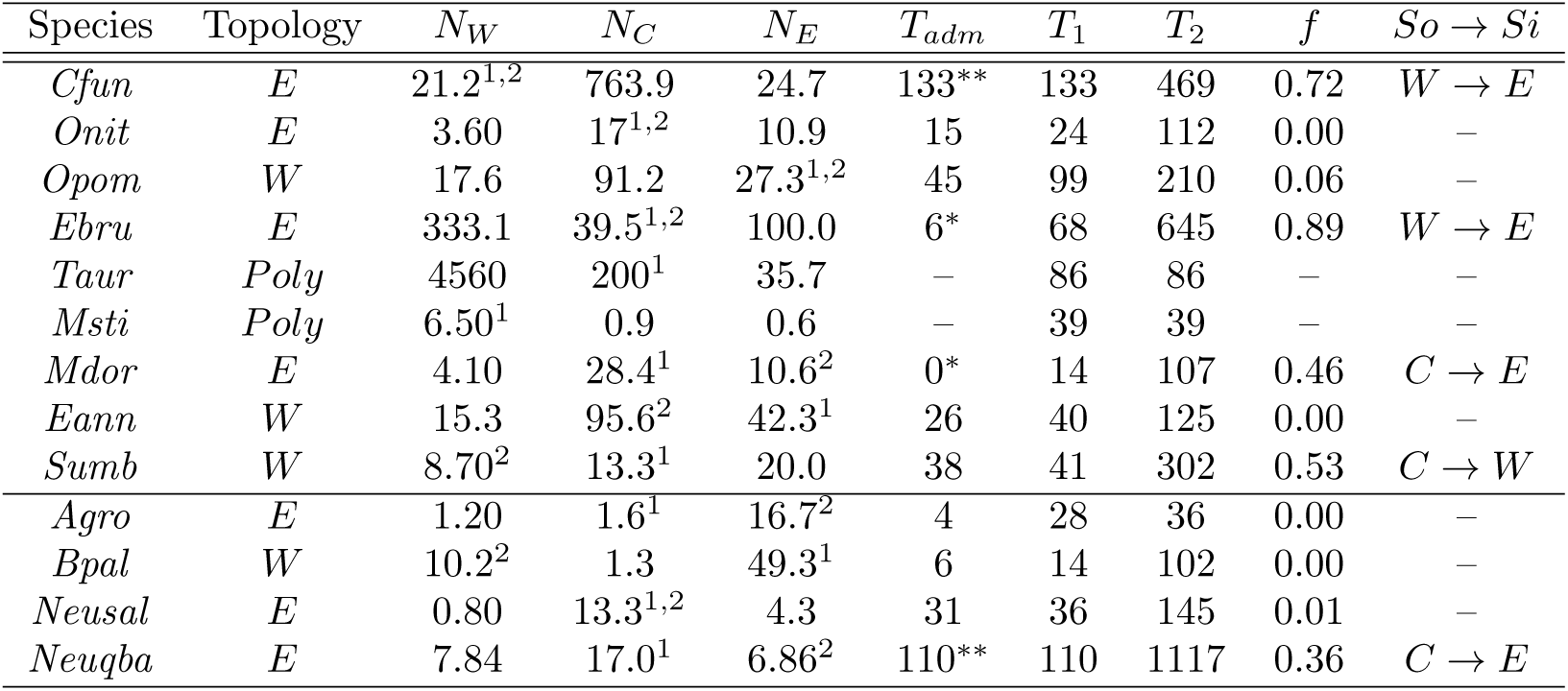
**Maximum likelihood estimates of demographic parameters and population topologies under the best supported model for each species**. Divergence (*T*_1_, *T*_2_) and admixture times (*T_adm_*), are given in thousands of years (ky), and effective population sizes (*N_W_*, *N_e_*, *N_E_*) are ×10^4^ individuals. Superscripts indicate the N_e_ shared by which ancestral population (younger = 1, older = 2). The superscript ^1,2^ indicates both ancestral populations share the same N_e_. For *T_adm_,* * indicates that the 95% confidence interval (C.I.) of the estimate for this parameter includes zero, while ** indicates that the 95% C.I. overlaps the 95% C.I. for *T*_1_. *So → Si* indicates the source and sink population for the admixture event. Species abbreviations: Cfun: *Cecidostiba fungosa,* Onit: *Ormyrus nitidulus,* Opom: *Ormyrus pomaceus,* Ebru: *Eurytoma brunniventris,* Taur: *Torymus auratus,* Mdor: *Megastigmus* dorsalis,Msti: *Megastigmus stigmatizans,* Eann: *Eupelmus annulatus,* Sumb: *Synergus umbraculus,* Agro: *Andricus grossulariae,* Bpal: *Biorhiza pallida,*, Neusal: *Pseudoneuroterus saliens,* Neuqba: Neuroterus quercusbaccarum.

| Species | Topology | $N_W$ | $N_C$ | $N_E$ | $T_{adm}$ | $T_1$ | $T_2$ | $f$ | $So \rightarrow Si$ |
| --- | --- | --- | --- | --- | --- | --- | --- | --- | --- |
| *Cfun* | *E* | 21.2<sup>1,2</sup> | 763.9 | 24.7 | 133** | 133 | 469 | 0.72 | $W \rightarrow E$ |
| *Onit* | *E* | 3.60 | 17<sup>1,2</sup> | 10.9 | 15 | 24 | 112 | 0.00 | – |
| *Opom* | *W* | 17.6 | 91.2 | 27.3<sup>1,2</sup> | 45 | 99 | 210 | 0.06 | – |
| *Ebru* | *E* | 333.1 | 39.5<sup>1,2</sup> | 100.0 | 6* | 68 | 645 | 0.89 | $W \rightarrow E$ |
| *Taur* | *Poly* | 4560 | 200<sup>1</sup> | 35.7 | – | 86 | 86 | – | – |
| *Msti* | *Poly* | 6.50<sup>1</sup> | 0.9 | 0.6 | – | 39 | 39 | – | – |
| *Mdor* | *E* | 4.10 | 28.4<sup>1</sup> | 10.6<sup>2</sup> | 0* | 14 | 107 | 0.46 | $C \rightarrow E$ |
| *Eann* | *W* | 15.3 | 95.6<sup>2</sup> | 42.3<sup>1</sup> | 26 | 40 | 125 | 0.00 | – |
| *Sumb* | *W* | 8.70<sup>2</sup> | 13.3<sup>1</sup> | 20.0 | 38 | 41 | 302 | 0.53 | $C \rightarrow W$ |
| *Agro* | *E* | 1.20 | 1.6<sup>1</sup> | 16.7<sup>2</sup> | 4 | 28 | 36 | 0.00 | – |
| *Bpal* | *W* | 10.2<sup>2</sup> | 1.3 | 49.3<sup>1</sup> | 6 | 14 | 102 | 0.00 | – |
| *Neusal* | *E* | 0.80 | 13.3<sup>1,2</sup> | 4.3 | 31 | 36 | 145 | 0.01 | – |
| *Neuqba* | *E* | 7.84 | 17.0<sup>1</sup> | 6.86<sup>2</sup> | 110** | 110 | 1117 | 0.36 | $C \rightarrow E$ |

### Species have expanded across the Western Palaearctic in different directions

A history of directional range expansion (*T*_2_ > *T*_1_) was supported for 11 of 13 species, while the two parasitoid species *Torymus auratus* and *Megastigmus stigmatizans* showed support for an unresolved polytomy (*T*_2_ = *T*_1_). Of the 11 species showing a directional signal, seven were inferred to have an Eastern origin (i.e. an (*E*, (*C, W*)) topology), including three of the four gallwasp species and four of the nine parasitoids (Table 1, Fig 2). In contrast, no species supported an (*C*, (*W, E*)) topology and a Western origin (i.e. a (*W*, (*C, E*)) topology) was supported for only four species (the gallwasp *Biorhiza pallida,* the inquiline gallwasp *Synergus umbraculus* and two parasitoids, Table 1). As expected, these same four species had lower mean divergence between the Centre and Eastern populations than between the West and either Centre and East (Fig S1). This incongruence in expansion histories argues against both strict co-dispersal and host tracking, but is compatible with ecological sorting and with random assembly.

**Fig 2.**
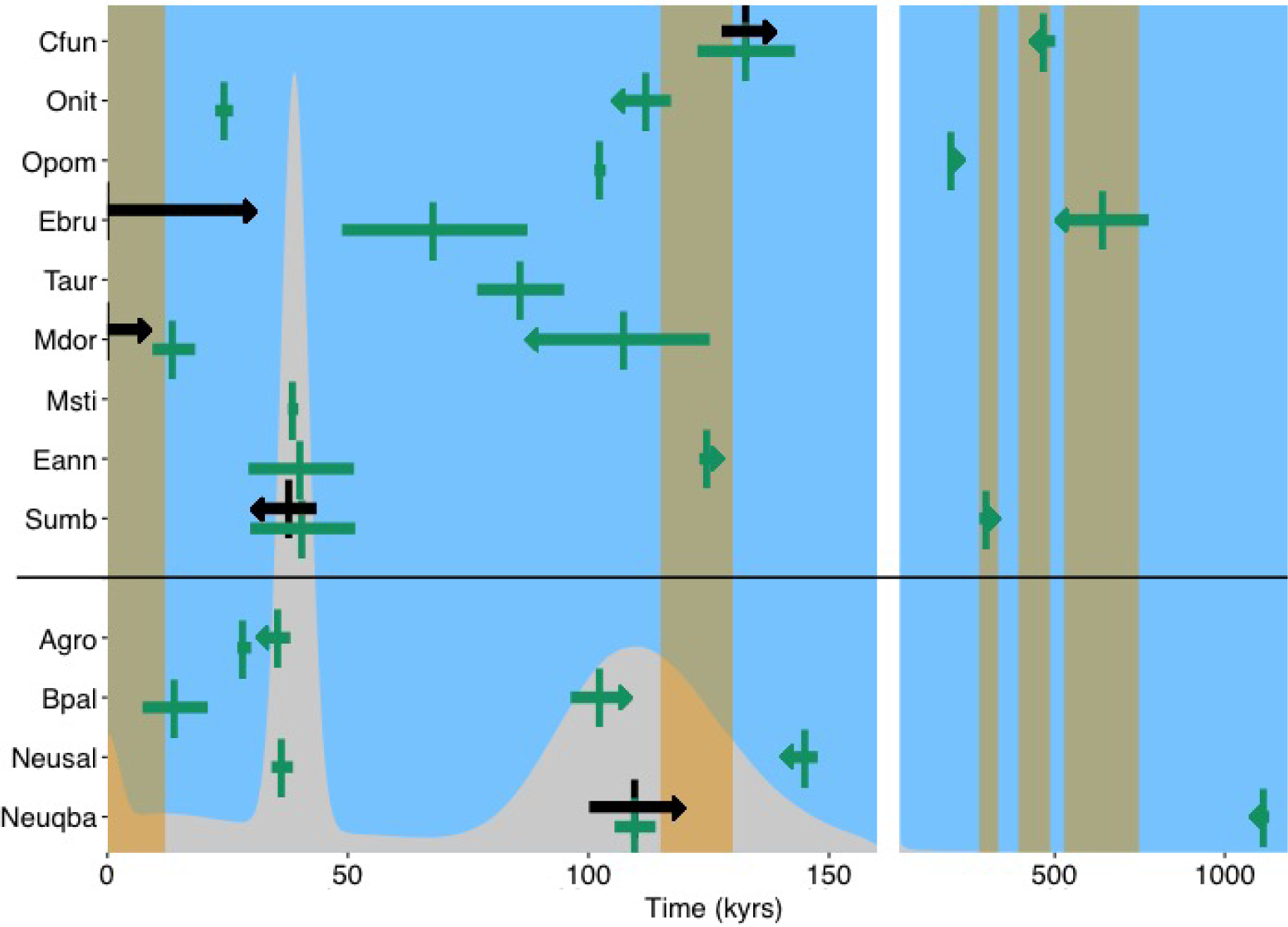
Community wide patterns of range expansion across the Western Palaearctic. Estimates for nine parasitoids and four gallwasp host species are shown at the top and bottom respectively (for species abbreviations see Table 1). Divergence times are shown in green, and comprise a point estimate flanked by 95% C.I.s. The arrow on the older divergence time for each species shows the direction of range expansion, e.g. a right-pointing arrow indicates expansion from West to East. Where supported in the best model, admixture times are shown in black. The arrow gives the direction of admixture, again comprising a point estimate flanked by 95% C.I.s. Orange vertical bars indicate interglacials. The best-fitting community-wide mixture distribution (see Methods) of divergence and admixture times is shown in grey. Note that the x-axis on the right-hand side of the figure is compressed relative to the left.

Admixture (*f* significantly > 0) between refugia was inferred for five species (indicated in black in Fig 2). All but one had an inferred Eastern origin with admixture back into the East either from the Centre or directly from the West. The exception was the inquiline *Synergus umbraculus,* which showed a (*W*, (*C, E*)) history with admixture from the Centre into the West, suggesting multiple waves of longitudinal dispersal in this species.

### The age of refugial populations differs across species but not guilds

The divergence times between refugia vary by over an order of magnitude across species, the oldest split (*T*_2_) ranging from ≈ 1100 thousand years ago (kya) in the gallwasp *Neuroterus quercusbaccarum* and 645kya in the parasitoid *Eurytoma brunniventris* to < 40kya in the gallwasp *Andricus grossulariae* and the parasitoid *Megastigmus stigmatizans* (Table 1). Given that a large number of species have non-overlapping confidence intervals (computed using a full parametric bootstrap, see Methods) for this split, we can confidently reject assembly via strict co-dispersal (Fig 2). Irrespective of whether we considered all species jointly or only those with a putative Eastern origin, the deepest population split (*T*_2_) was not consistently older in gallwasp hosts than in their parasitoid enemies (Fig 2), arguing against delayed host tracking and widespread enemy escape. We stress that our comparison of population divergence times across species does not rely on any absolute calibration but only assumes i) knowledge of the generation times of each species (which we have, see Methods, [16]) and equal mutation rate across species (which seems reasonable) (see Table S3 for uncalibrated parameter estimates).

### Community-wide expansion pulses coincide with warm periods in the Pleistocene

Our rejection of the co-dispersal and host pursuit models raises the question of whether the high observed diversity of species’ demographic histories is nonetheless structured, or is compatible with a neutral assembly model. We address the first question by asking how many divergence events are required to explain the demographic history of this community? Considering only the oldest split in each species (*T*_2_), we explored models of co-divergence (using a discretized grid of the marginal support for *T*_2_, see Methods). The most parsimonious model of co-divergence contained two clusters of species with indistinguishably similar divergence times *(Andricus grossulariae* and *Megastigmus stigmatizans* at 37 kya and *Torymus auratus, Megastigmus dorsalis* and *Biorhiza pallida* at 102 kya) and eight species that each diverged independently of all others (Δ*lnL* = 3.15, 3 d.f., *p* = 0.0979, Table S4).

A second, and as we would argue, more meaningful question is whether the assembly of the oak gall community is compatible with an altogether random process. In other words, can we reject an ecologically neutral null model that views species as equivalent? To test this we fitted continuous distributions to the community-wide set of divergence (*T*_1_ and *T*_2_) and admixture (*T_adm_*) times. This was done in a new comparative framework (see Methods) that allows us to include uncertainty in species-specific estimates. Arguably [10], the simplest ecologically neutral model would view the timing of divergence and admixture between refugia as a simple waiting process (time parameters are drawn from an exponential distribution) or assume a single unimodal distribution (e.g., a log normal). We were able to reject both of these null models of community assembly in favor of a more complex history involving several continuous multi-species divergence pulses (Δ*lnL* = 2.61, 3 d.f., *p* = 0.156). The most parsimonious model assumes that, community-wide, the times of between-refuge divergence and admixture are given by a mixture distribution consisting of a single exponential and three log normal distributions (Fig 2) with modes of 10, 39 & 110 kya and a combined probability mass of 0.96.

The fact that these coincide approximately with warm periods in the late Pleistocene (corresponding to the beginning of the current Holocene interglacial, a Dansgaard-Oeschger interstadial event, and the Eemian interglacial) suggests that, within the oak gall community, expansions into and admixture between refugia are associated with periods of geographic expansion of the distributions of oaks and their associated gall communities – a process confirmed for the Eemian interglacial by fossils for both gallwasps and parasitoids [23]

## Discussion

Understanding how complex natural communities have assembled requires a retrospective approach, and inference of the population histories of component species. Previous comparative analyses of the phylogeographic history of trophically-linked species have either been limited in scope by the small numbers of species compared (e.g. [13, 24]) and/or were based on small numbers of loci and achieved low resolution of the demographic histories of individual species [16]. Here we use whole genome data for small samples of individuals to make detailed, continental-scale reconstructions of the Pleistocene histories of 13 members of a widespread insect community: oak gallwasps and their associated parasitoid enemies.

Our systematic comparison of population histories within and between trophic levels unveils considerable complexity in the assembly of the oak gall community. This can neither be captured by classic models of deterministic community assembly which predict a small number of co-divergence events [25], nor an ecologically neutral process which predicts a simple community-wide distribution of divergence times [10]

Members of the oak gall community differ in both the directionality and timing of their longitudinal expansion into Europe, without any consistent difference between gallwasp hosts and their parasitoids. We can therefore confidently reject both co-divergence and enemy escape, two contrasting and paradigmatic models of community assembly. Moreover, the fact that we identify a minimum of nine distinct co-divergence events for the oldest split implies that we can also rule out more complex assembly scenarios that involve a small number of co-divergence events, e.g., a history where all species co-diverge with at least one other. Our comparative analysis also allows us to reject an ecologically neutral null model of random community assembly. This model views species as exchangeable and assumes that divergence and admixture events between refugia happen at a single community-wide rate [10, 9] and independently of past climatic events.

### Ecological sorting and host range

Our finding that species differ both in the timing and directionality of range expansions shows that the stability of interactions between them varies in space. For example, we infer that the seven species showing an out of the East population history have shared the Eastern refuge throughout their history. In contrast, interactions in the Centre and the West between any two of the same set of species are only possible when both have reached these refugia. Because divergence times in both guilds vary by an order of magnitude there must have been considerable turnover in species interactions. For example, *Eurytoma brunniventris*, which has the most ancient out of the East parasitoid population history, can only have exploited hosts whose expansion histories into the Centre are at least as early (such as *Neuroterus quercusbaccarum*). Interactions in the Centre with later expanding hosts, such as *Andricus grossulariae,* could only be restored when these, too, arrived from the East. The more out of step expansion times become, the more interactions must have been disrupted and the lower the potential for tight coevolution. Such turnover in trophic relationships may also explain why this and other communities centered around temperate insect herbivores are dominated by relatively generalist species, which attack a range of hosts within the community [26, 7].

An alternative explanation for the incongruence in timing we observe across species is that the genetic signatures of older demographic events have been erased by extensive subsequent admixture in some species, but not in others. While such potential loss of signal is a general limitation of demographic inference and cannot be ruled out, it seems an unlikely explanation for our general finding of incongruence between species, given the absence of admixture signals in most of them.

A species’ potential to expand its range is a function of its ecology and life history. A recent study identified life-history strategy as the only correlate of genetic diversity: animals with more rapid life histories and larger numbers of offspring had higher genetic diversity [20]. While this and other comparative studies of genetic diversity [21, 27] have ignored demography, comparing species with respect to their demographic histories makes it possible (at least in principle) to test for assembly rules. For example, if ecological sorting plays an important role in community assembly, more generalist and/or widespread parasitoids should have older histories than specialists. We do indeed find that *T*_2_ and ancestral *N_e_* are positively correlated with both host range and abundance, which is compatible with this idea (Fig S2). However, confirming this relationship will require larger samples of species.

### Eastern origins and Westerly winds

Most, but not all members of the oak gall community have an Eastern origin. This is not surprising given the pattern of decreasing genetic diversity from East to West found by previous studies of this community [12, 13, 24] and Western Palaeartic taxa more generally [28, 29] and the inference of regional gallwasp community diversification ca. 10 mya in Anatolia/Iran [15]. While only a few species in the oak gall community show evidence for admixture, it is intriguing that i) admixture appears most substantial in the three species with the oldest histories: *Neuroterus quercusbaccarum, Cecidostiba fungosa* and *Eurytoma brunniventris* (Table 1) and ii) it occurred predominantly from the West into the East, that is, in the opposite direction of population divergence. We conducted a simple simulation study to confirm that this directional signal is not an artifact of our modelling approach, i.e., admixture back into the source population is not easier to detect than in the reverse direction (see Methods). Given that both chalcid parasitoids and cynipids have been observed in aerial samples taken 200m from the ground [30] and the prevailing winds in the Western Palaearctic are from the West, it seems plausible that gene flow into the East is simply easier and therefore more likely than in the reverse direction.

### Limitations and potential confounding factors

While the demographic models we consider are necessarily simplifications of the truth, they capture key population level processes that have been largely ignored by previous comparative analyses. First, many comparative phylogeographic studies have focused on divergence between only pairs of populations [16, 31] Second, analyses of divergence and admixture between multiple populations have, for the sake of tractability, often assumed equal *N_e_* among populations [32, 33] which can bias estimates of divergence and admixture. Our finding that N_e_ varies by up to two orders of magnitude among populations within species (Table 1) highlights the importance of modeling N_e_ parameters explicitly [34, 35]. While this study’s reanalysis of *B. pallida* inferred little admixture when accounting for *N_e_* differences between refugia, a previous analysis based on a single sample per population [32] suggested substantial gene flow from East to West under a model in which ancestral *N_e_* was fixed across all populations.

To make robust inferences about community assembly, it is imperative to minimize biases that might affect comparability among species. We have sequenced species to the same coverage whenever possible and have used the same bioinformatic pipeline and comparable block lengths (in terms of diversity) for all species. To test whether selective constraint is likely to differ drastically among species or trophic levels (which could lead to systematic biases in demographic estimates), we compared diversity across all sites to that at synonymous coding sites and found no difference between between hosts and parasitoids (Suppl. Info 2). Finally, our parametric bootstrap replicates which were simulated with recombination show that intra-locus recombination rates (which may differ between species) had minimal effects on model and parameter estimates.

### A new comparative phylogeography

Previous comparative studies have been unable to build such a nuanced picture of community assembly in this or any other system due to limitations in both available data and analytical approaches [16, 36, 37] (although new methods have begun to emerge [38, 39]). In particular, we conducted a previous analysis of the oak gallwasp community based solely on mitochondrial data sampled for the same set of refugia and an overlapping set of taxa. In contrast to the present study, this previous analysis had very limited power to resolve even simple demographic histories with or without gene flow [16].

We now have both the inference tools and the data necessary to resolve divergence and admixture relationships among multiple populations and across many trophically linked species. Given the resolution of whole genome data, it becomes hard to justify the logic of testing for strict co-divergence [40], if this implies identical divergence times across species. Three recent comparative phylogeographic analyses, each based on thousands of ddRAD loci, avoided using approaches that test for strict co-divergence, but instead compared parameter estimates post hoc among species [41, 42, 43]. For example, Oswald et al. [42] found that six pairs of co-distributed birds shared the same mode of divergence (isolation with asymmetric migration) but diverged asynchronously. Our approach replaces informal post hoc comparisons with a new formal comparative framework that both incorporates uncertainty in species-specific parameter estimates and characterises community-wide distributions of divergence times.

### Outlook

Being able to efficiently reconstruct intraspecific histories from genome-scale data across multiple species means that one can infer the tempo and mode of community assembly across space and through time. The composite likelihood framework we use applies to a wide range of demographic histories [44, 24] and, as we have shown here, lends itself to comparative analyses. Given sufficient replication in terms of species, such analyses open the door to answering a range of fundamental questions in evolution and ecology. For example, an interesting avenue of future research will be to explicitly include selection in comparative analyses. In particular, it will be fascinating to ask if and how range shifts and admixture events are accompanied by selection on genes that may be involved in host-parasitoid interactions. It will soon be possible to use whole genome data to compare not only the demographic histories of interacting species but also the signatures of selective sweeps and their targets in the genome. In practice, such more detailed comparisons will require both further work on inference methods and more complete datasets, including more contiguous (and functionally annotated) reference genomes and larger samples of individuals and taxa.

## Materials and Methods

### Samples and sequencing

#### Sample processing

We generated WGS data for three species of oak gallwasp hosts, eight parasitoids and one inquiline. We also reanalysed WGS data generated for a pilot study for *B. pallida* another gallwasp species [32] (ENA Sequence Read Archive: http://www.ebi.ac.uk/ena/data/view/ERP002280). For each species we sampled a minimum of two male individuals from each of three refugial populations spanning the Western Palaearctic: Iberia (West), the Balkans (Centre) and Iran (East) (Table S2). Only a single Eastern sample was available for two species: *Ormyrus pomaceus* and *Biorhiza pallida*.

DNA was extracted from (haploid) male individuals using the Qiagen DNeasy kit. Nextera libraries were generated for each individual specimen and sequenced on a HiSeq 2000 by Edinburgh Genomics. Raw reads are deposited in the Short Read Archive (Project code: PRJEB15172). Median average coverage per species ranged from 3.81 to 6.59 for the parasitoids and 2.94 to 9.33 for the gallwasps. The inquiline gallwasp, *Synergus umbraculus* was sequenced to higher coverage (median 34.1) because its genome size was initially over-estimated.

#### *De novo* genome assembly

Reads were quality controlled and adapter trimmed using *trimmomatic* [M1], followed by *cutadapt* [M2] to remove remaining *Nextera* transposase sequences. Results were inspected using *FastQC* [M3]. For each species, data were combined across all individuals and a reference genome assembly generated using *SPAdes* [M4] or *MaSuRCA* [M5] (Table S1). We used *RepeatScout* [M6] and *RepeatMasker* [M7] to *de novo* predict repeat regions. Contigs that *Blast* [M8] matched putative contaminant (bacteria, fungi) or mitochondrial sequence were removed from further analysis.

#### Variant calling

Reads per individual were mapped back to each reference genome using *BWA* (0.7.15-r1140) [M9] and duplicates marked using *picard* http://broadinstitute.github.io/picard/(V2.9.0) MarkDuplicates. Variant calling was performed with *freebayes* http://github.com/ekg/freebayes(v1.1.0-3-g961e5f3) with minimum alternate allele count (-C) of 1, minimum base quality (-q) of 10, minimum mapping quality (-q) of 20. Variants were then filtered for a minimum quality of 20 and decomposed into allelic primitives. Remaining single nucleotide polymorphisms (SNPs) were retained for further analyses.

#### Generating blockwise data

The CallableLoci walker of GATK (v3.4) [M10] was used to identify regions covered in all individuals and with a base quality > 10 and a mapping quality > 20 in the mark-duplicated bam files (these filters are the same used for SNP calling in *freebayes*). Blocks of a fixed length l of callable sites were generated for each species using custom scripts (deposited at *http://github.com/KLohse*). For parasitoids, we only included sites that passed the CallableLoci filter in all six individuals. Given the overall lower sequencing depth for the gallwasps, we relaxed this requirement in three of the four gallwasp hosts *(Andricus grossulariae, Biorhiza pallida,* & *Pseudoneuroterus saliens*) and partitioned the data into blocks using the CallableLoci filter independently for each subsample of *n* = 3 (see below).

Since our inference framework assumes no recombination within blocks, we partitioned the data into short blocks (and performed a simulation based check for potential biases). To ensure equal information content and to minimize differences in bias due to recombination within blocks across species, we generated blocks with a fixed average number of 1.5 pairwise differences. In other words, the physical length of blocks *l* was chosen to be inversely proportional to the average pairwise per site diversity *π* (post filtering) of a species and so differed between taxa.

Blocks with a physical span (pre-filtering length) of > 2*l* or containing more than five ‘None’ sites were removed. Likewise, short contigs (< 2 × *l*) were ignored. These filtering steps resulted in datasets of between 55,909 and 635,041 blocks per species. For the three gallwasp species where each subsample was filtered independently, average block number varied between 19,303 and 86,588 blocks per species.

### Fitting demographic models

#### Computing model likelihoods

Models were fitted to block-wise data using an analytic coalescent framework [19]. Briefly, the probability of observing a particular mutational configuration in a sequence block can be expressed as a higher order derivative of the generating function of genealogical branch lengths. This calculation assumes an infinite sites mutation model (which is reasonable given the low per site diversity). It is straightforward to calculate the support (i.e. the logarithm of the likelihood, *lnL*) of a particular model, given the corresponding generating function and a table of counts of observed block-wise mutational configurations.

We used an automation [18] previously implemented in *Mathematica* [M11] to obtain the generating functions for the set of 48 models. Since a solution for the GF for the full sample of six individuals (two per population) is intractable, we computed the following composite likelihood:

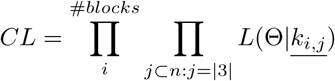

where <u>*k_i,j_*</u> is the count of the mutations defined by the three unrooted branches for block *i* and triplet *j*. In other words, the composite likelihood *(CL)* is a product over contributions from all (14 for a sample of size *n* = 6 we consider) possible (unrooted) subsamples of triplets and all blocks. This calculation scales to arbitrarily large samples of individuals as the GF expressions for each triplet are small. It also makes use of the phase information contained in the haploid data. Unlike previous implementations (i.e. [32]) there is no requirement for an outgroup. Analogous composite likelihood strategies based on subsamples have been used in phylogenetic network analysis [M12] and to infer divergence and continuous gene flow [M13, M14]. We maximized *lnCL* for all parameters (*N_E_, N_e_, N_W_*, *T*_1_,*T*_2_,*T_adm_, f*) included in the best supported model using a Nelder-Mead simplex algorithm implemented in the *Mathematica* function *NMaximize.* Configurations involving more than *k_max_* = 2 mutations on any particular branch were combined in the *CL* calculation. The method differs from other recent approaches for inferring reticulate evolutionary histories in that it allows *N_e_* to differ between populations (cf. [M12]) and makes use of linkage among sites (cf. TreeMix [M15]).

#### Parametric bootstrap

Maximizing the *CL* across blocks and subsamples ignores linkage between blocks but gives an unbiased point estimate of parameters. To measure the uncertainty in demographic parameter estimates we conducted a full parametric bootstrap: for each species, we simulated 100 replicate datasets in *msprime* [M16] under the full ancestral recombination graph and the best supported model (and the MLEs under that model). Recombination rates per base pair (*ρ*) were inferred for each species using the two-locus generating function outlined in [19]). This inference was based on two Spanish individuals from each species (see File S1). Each bootstrap replicate had the same total sequence length as the real dataset (after filtering). We assumed that blocks in the bootstrap replicates were immediately adjacent to one another which is conservative given the filtering pipeline used to delimit blocks in the real data. For the sake of computational efficiency, each simulation replicate was divided into 20 equally sized chunks/chromosomes. 95% confidence intervals (C.I.) were obtained as two standard deviations of estimates across bootstrap replicates.

#### Identifying co-divergence clusters

To assess whether the oldest divergence times *T*_2_ differed significantly between species and trophic levels, we implemented a hierarchical clustering procedure. For each of the eight species that overlapped in 95% C.I. for *T*_2_ with any other species, we evaluated the marginal *lnCL* along a discretized grid (maximising *lnCL* across all other parameters). We used the standard deviation in T2 estimates across parametric bootstrap replicates to rescale these marginal *lnCL.* These rescaled *InCL* were treated as an approximation of the true marginal support *(lnL)* for *T*_2_.

To test whether divergence times differed between species, we computed the support for models in which divergence times are shared by subsets of taxa. Essentially, this phrases the test for clustered divergence times as a model simplification problem: for example, the MLE for a co-divergence event involving two or more species is given by the maximum of the sum of their respective marginal *lnL.* The associated reduction in model support is measured by *ΔlnL* relative to the full model, i.e., allowing species to have unique divergence times. We identified the most parsimonious model as the model with the fewest distinct T_2_ clusters (assuming *ΔlnL* has a χ^2^ distribution) and examined all possible combinations of co-diverged species.

#### Characterizing the community-wide distribution of divergence and admixture times

We fit a series of distributions of increasing complexity to the inferred *T* parameters for all species. We assumed that divergence and admixture times across species are given by some community-wide function *g*(*t*; λ). The likelihood of λ, the community-wide rate of population divergence (or set of parameters in the case of more complex functions) is:

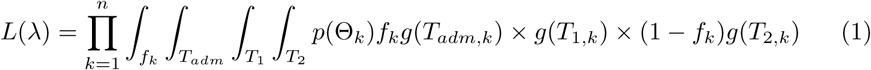

where Θ_*k*_ is the set of time parameters and the admixture traction tor the kth species and *p*(Θ_*k*_) is the (normalized) likelihood of Θ_*k*_. Since computing this likelihood directly from the data is intractable, we used a discretized grid of likely *T* and *f* values (i.e. replacing the integrals in (1) by sums). Grids were centered around the MLE and bounded by the 95 % C.I. of parameters. In each dimension, we evaluated 5 points. For

computational tractability, we fixed *θ* and the *N_e_* scalars at their MLEs. We initially fitted exponential and log normal distributions, followed by increasingly complex mixture distributions of a single exponential and several log normals. Models were fitted sequentially: as we included additional mixture contributions, we fixed the parameters and mixture weights of the lowest weighted component of the previous distribution. We identified the most parsimonious model as the simplest model (i.e. with the fewest parameters) which did not give a significant reduction in *lnL*.

## Acknowledgments

We thank Richard Harrison, James Nicholls, Sarah White and Lisa Cooper for help in the lab and Edinburgh Genomics for sequencing. Many thanks to Stuart Baird, Alex Hayward and Mike Hickerson for discussions throughout the project and to Ally Phillimore for a critical manuscript review. We are grateful to Rob Ness for advice on the bioinformatics, to Pablo Fuentes-Utrilla, György Csóka, George Melika and Majide Tavakoli for contributing specimens, to Richard R. Askew for help with identification and William Walton for collating the ecological data. This work was supported by a standard grant from the Natural Environmental Research Council (NERC) UK to GNS and KL (NE/J010499/1). KL is supported by a NERC Independent Research fellowship (NE/L011522/1).

## Author contributions

GNS & KL conceived the study, LB conducted analyses, JK developed & implemented the bioinformatic pipeline, KL developed the analytic framework with contributions from LB. LB, KL & GNS wrote the manuscript.

## Supporting Information

**Table S1.** **Assembly summaries**. N50: a weighted median statistic such that 50% of the entire assembly is contained in contigs or scaffolds equal to or larger than this value. CEGMA (Core Eukaryotic Genes Mapping Approach): % complete or partial presence of a core set of eukaryotic genes. See Table 1 for species abbreviations.

| Species | Assembler | N50 | Assembly size<br>> 200 bases | Number of contigs<br>> 200 bases | Cegma<br>% complete | Cegma<br>% partial | Median average<br>coverage per<br>individual |
| --- | --- | --- | --- | --- | --- | --- | --- |
| Cfun | SPAdes | 87417 | 182108758 | 18121 | 93.55 | 95.97 | 6.59 |
| Onit | SPAdes | 22984 | 259884369 | 37082 | 95.16 | 99.19 | 5.69 |
| Opom | SPAdes | 31593 | 263296421 | 43391 | 93.15 | 97.98 | 5.97 |
| Ebru | SPAdes | 3829 | 393847642 | 200270 | 90.73 | 98.79 | 5.74 |
| Taur | SPAdes | 10570 | 397870892 | 100431 | 91.53 | 97.18 | 5.14 |
| Msti | MaSuRCA | 9143 | 577876374 | 182922 | 93.55 | 97.98 | 3.81 |
| Mdor | MaSuRCA | 18748 | 589959111 | 148702 | 95.16 | 97.58 | 4.34 |
| Eann | SPAdes | 28524 | 314955021 | 48024 | 92.74 | 97.58 | 6.23 |
| Sumb | MaSuRCA | 49302 | 235334141 | 20256 | 95.97 | 98.79 | 34.1 |
| Bpal | CLC | 1075 | 805102378 | 1163314 | 32.26 | 67.74 | 9.34 |
| Neuqba | CLC | 1664 | 2569683019 | 2812183 | 22.58 | 60.08 | 3.17 |
| Neusal | CLC | 970 | 2060851172 | 3378461 | 45.56 | 80.65 | 2.94 |
| Andgro | CLC | 1511 | 1877197301 | 2221463 | 37.10 | 72.58 | 5.69 |

### Supp. Info 1 Comparing genetic diversity within and divergence between refugia across the community

Although genetic diversity estimates for species in the oak gallwasp community lie within the range previously found for a taxonomically diverse set of arthropods [20], they vary by an order of magnitude in both the herbivore and parasitoid guilds (Fig S1). While genetic diversity differs between refugia, we find no general community-wide pattern of declining within-refuge genetic diversity across species as expected for a simple range expansion from East to West (which predicts *π_E_* ≤ *π_C_* ≤ *π_W_*, as in the parasitoid *Megastigmus dorsalis* and the gallwasp *Neuroterus quercusbaccarum*) or from West to East (which predicts the converse, as shown by the parasitoid *Torymus auratus*). Across all species, diversity is slightly lower in the West (0.00230), than in the Center and East (0.00288 and 0.00284 respectively), with no significant differences between refugia (Fig S1).

Divergence (as measured by *d_XY_* [S1]) between refugial populations exceeds diversity within them, confirming that species in this community are indeed structured into refugia (Fig S1). For eight out of 13 species, mean divergence between the East and either the West or the Centre populations (*d_EW_* and *d_EC_*) exceeds divergence between the West and Centre (*d_WC_*), consistent with range expansion from the East (i.e. a population tree topology ((*E*, (*C,W*))). Further, for a history of strict divergence without gene flow and topology (*E*, (*C, W*)), we expect *d_EW_* = *d_EC_* (this is the analog of the D statistic [S2] for unrooted samples). In contrast to this prediction, several species show asymmetries in pairwise *d_XY_*, a pattern consistent with gene flow between refugia after their initial divergence.

**Fig S1.**
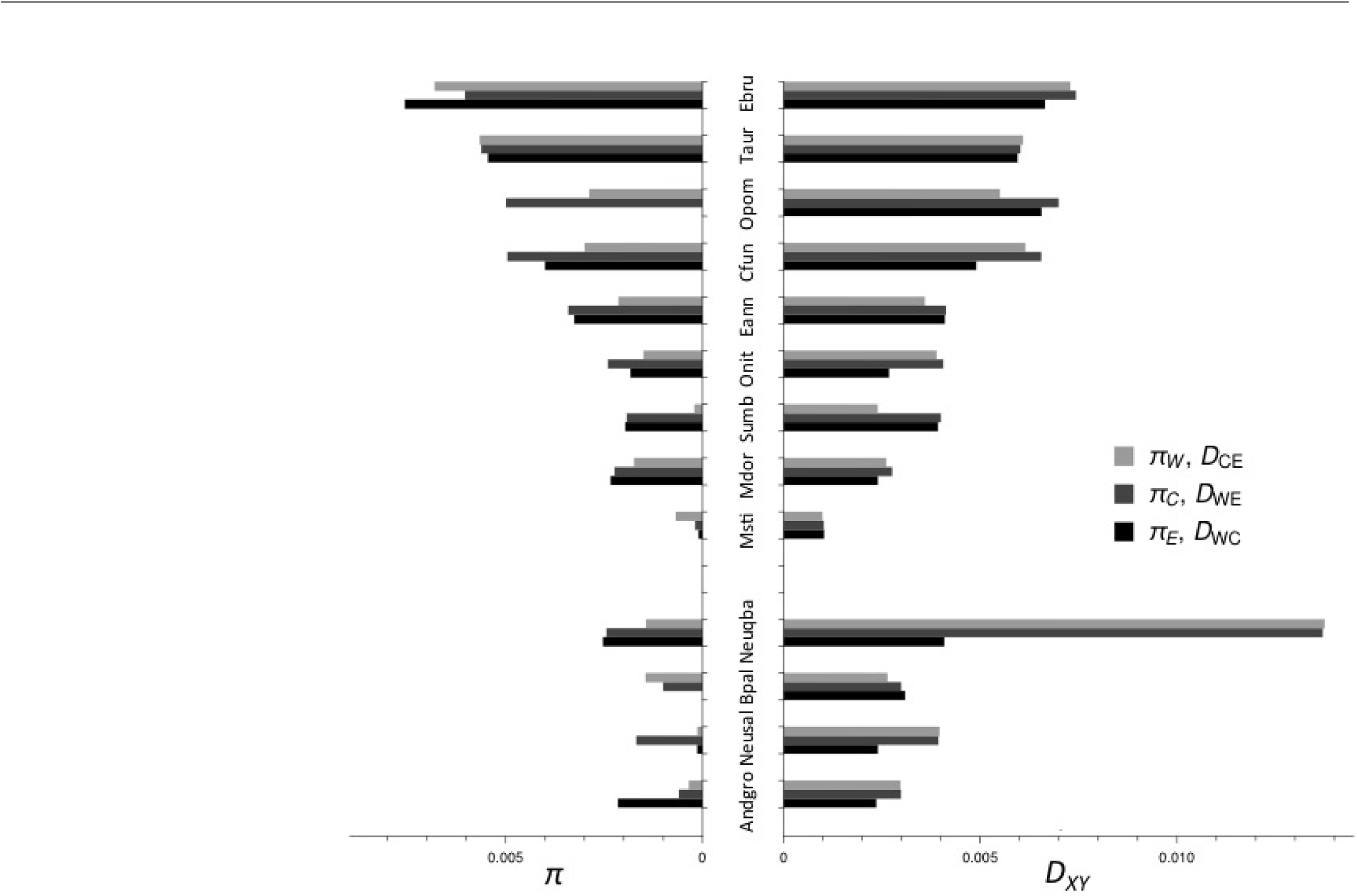
Genetic diversity within & divergence between refugia for 13 species from the oak gall community. Parasitoids above and gallwasp hosts below. Bars on the left show pairwise per base nucleotide diversity (*π*) for individuals sampled in the Western (W), Central (C) & Eastern (E) refugium. Bars on the right show pairwise nucleotide divergence (*d_XY_*) between refugia. Note that *π_E_* is missing for *Opom* and *Bpal* because only a single individual was available from the East. See Table 1 for species abbreviations.

**Fig S2.**
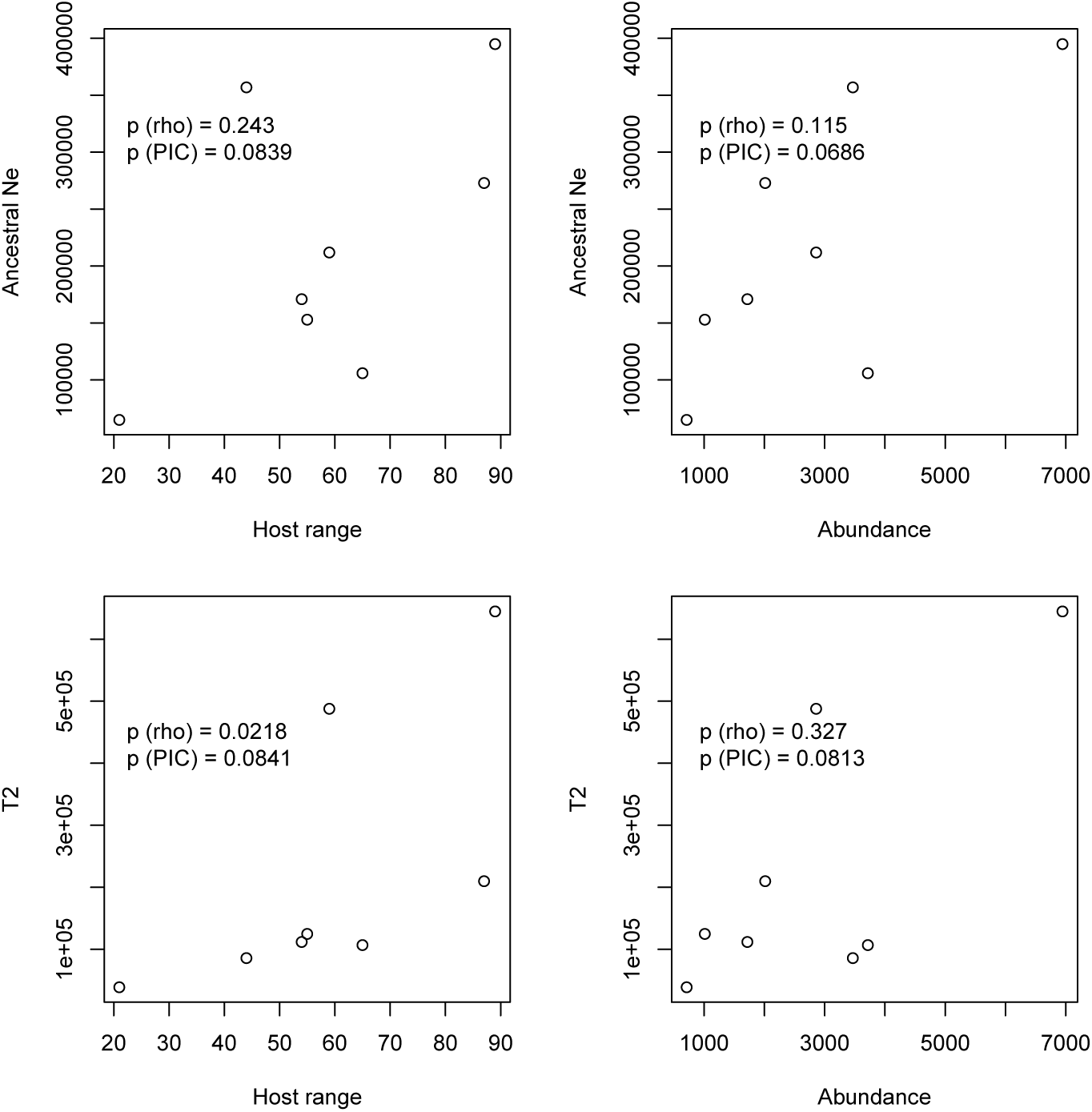
Ecological correlates of parasitoid demographic parameters. Host range was calculated as the number of gallwasp host species each species has been successfully reared from. Abundance was calculated as the number of individuals of each species reared in total from all hosts combined. Both measures were taken from [S3], itself a compilation of published sources and the authors’ own rearing data. Ancestral *N_e_* (of older ancestor) and *T*_2_ are the ML parameter estimates from the best-fit model for each species (Table 1). P values cited are for Spearman’s rank correlations and phylogenetic independent contrasts (sister group comparisons only) calculated using the R package ‘caper’ [S4].

### Suppl. Info 2 Generating constraint data & testing for selective constraint

SNP-converted fasta sequences were created for each individual using the SelectVariants and FastaAlternateReferenceMaker walkers of GATK and VCF files. Potential coding regions were identified in each genome assembly by *BlastX* using the gene set of the chalcid *Nasonia vitripennis* (http://arthropods.eugenes.org/genes2/nasonia/genes/) for parasitoids and an oak gallwasp gene set from *Biorhiza pallida* [32] for gallwasps. These matching regions were expanded by up to 10 000 bp in flanking sequence and then refined into coding sequence using *Exonerate* [S5] and the corresponding highest-scoring *BlastX* protein to define exons. The resulting coding sequence coordinates were used to excise sequences from the per individual fasta sequences with *bedtools* [S6] and custom scripts. Multiple sequence alignments were created for all individuals per species with *translatorX* [S7]. Alignments containing internal stop codons were removed. Four-fold degenerate sites and polymorphisms were predicted for each alignment with polydNdS of the *libsequence* package [S8]; alignments containing polydNdS-predicted internal stop codons were removed.

Given that assembly quality and genome composition varied among species, we tested whether the average level of constraint due to background selection differed between species. Average constraint was measured as the ratio of mean within population diversity at all sites (*π_a11_*) relative to the diversity at synonymous (four-fold degenerate) sites (n_s_) in single copy coding genes, which we assumed to evolve neutrally [S9]. As expected, *π_all_* was smaller than *π_s_* for the majority of species: *π_all_*/*π_s_* ranged from 0.58 to 1.36. Importantly, however, selective constraint as measured by *π_all_*/*π_s_* did not differ between trophic level (Mann-Whitney test, U = 21.5, p = 0.643) and was not correlated with assembly quality (as measured by N_50_, Spearman’s rho = 0.0165, p = 0.957) or size (Spearman’s rho = 0.206, p = 0.499). We therefore conducted all analyses without species-specific constraints.

**Table S2.** Specimens used for genome sequencing Sampling location for individual wasps used for genome sequencing (see Table 1 for species abbreviations). See [32] for *Biorhiza pallida.* Top table contains gallwasp species, bottom table contains parasitoid species.

| Species | Individual | Locality country | Locality region | Locality name | Latitude | Longitude |
| --- | --- | --- | --- | --- | --- | --- |
| Neuqba | Neuqba139 | Spain | Cádiz | Benamahoma | 36.77 | -5.47 |
|  | Neuqba230 | Spain | Cádiz | Benamahoma/Grazalema | 36.76 | -5.44 |
|  | Neuqba64 | Hungary | - | Mátrafüred | 47.83 | 19.97 |
|  | Neuqba65 | Hungary | - | Mátrafüred | 47.83 | 19.97 |
|  | Neuqba237 | Iran | Lorestan | Kaka-Sharaf | 33.38 | 48.46 |
|  | Neuqba236 | Iran | Lorestan | Kaka-Sharaf | 33.38 | 48.46 |
| Neusal | Neusal153 | Spain | Cáceres | La Aliseda | 40.33 | -5.38 |
|  | Neusal155 | Spain | Madrid | El Pardo | 40.53 | -3.77 |
|  | Neusal32 | Hungary | - | Szentkut | 47.98 | 19.78 |
|  | Neusal12 | Hungary | - | Mátrafüred | 47.83 | 19.97 |
|  | Neusal156 | Iran | Lorestan | Kaka-Sharaf | 33.38 | 48.46 |
|  | Neusal158 | Iran | Lorestan | Taf-Tfina | 33.30 | 48.45 |
| Andgro | Andgro421 | Spain | Ávila | - | 40.60 | -4.48 |
|  | Andgro426 | Spain | Ávila | - | 40.60 | -4.48 |
|  | Andgro319 | Hungary | - | Godollo | 47.60 | 19.38 |
|  | Andgro317 | Hungary | - | Godollo | 47.60 | 19.38 |
|  | Andgro425 | Iran | Lorestan | Pirshamse | 30.26 | 51.35 |
|  | Andgro422 | Iran | Lorestan | Ghelaei | 33.78 | 47.93 |
| Mdor | Mdor1348 | Spain | Salamanca | Horcajo | 40.64 | -5.41 |
|  | Mdor1457 | Spain | Madrid | El Escorial | 40.58 | -4.13 |
|  | Mdor1350 | Hungary | - | Isaszeg | 47.53 | 19.40 |
|  | Mdor1448 | Hungary | - | Godollo | 47.60 | 19.38 |
|  | Mdor2932 | Iran | Lorestan | Ghelaie | 33.78 | 47.93 |
|  | Mdor2934 | Iran | Kordestan | Marivan | 35.52 | 46.17 |
| Msti | Msti0395 | Spain | Salamanca | Horcajo | 40.64 | -5.41 |
|  | Msti0314 | Spain | Madrid | El Escorial | 40.58 | -4.13 |
|  | Msti0553 | Hungary | - | Sástó | 47.84 | 19.96 |
|  | Msti0548 | Hungary | - | Sástó | 47.84 | 19.96 |
|  | Msti0904 | Iran | - | Golestan | 36.84 | 54.44 |
|  | Msti0498 | Iran | - | Shena | 33.55 | 48.08 |
| Cfun | Cfun135 | Spain | - | Puerto de Villatoro | 40.55 | -5.17 |
|  | Cfun142 | Spain | Madrid | El Escorial | 40.58 | -4.13 |
|  | Cfun83 | Hungary | - | Sopron | 47.68 | 16.60 |
|  | Cfun71 | Hungary | - | Mátrafüred | 47.83 | 19.97 |
|  | Cfun86 | Iran | Kermanshah | Gahvareh | 34.35 | 46.42 |
|  | Cfun3549 | Iran | Lorestan | Taff | 33.30 | 48.45 |
| Opom | Opom344 | Spain | Cáceres | Aldeanueva del Camino | 40.26 | -5.93 |
|  | Opom673 | Spain | Ávila | La Cañada | 40.60 | -4.48 |
|  | Opom688 | Hungary | - | Mátrafüred | 47.83 | 19.97 |
|  | Opom687 | Hungary | - | Mátrafüred | 47.83 | 19.97 |
|  | OrmPom | Iran | Azerbaijan West | Piranshahr | 36.70 | 45.14 |
| Onit | Onit386 | Spain | Avila | Lanzahita | 40.20 | -4.93 |
|  | Onit350 | Spain | Madrid | Cotos de Monterey | 40.80 | -3.61 |
|  | Onit1632 | Hungary | - | Isaszeg | 47.53 | 19.40 |
|  | Onit1633 | Hungary | - | Godollo | 47.6 | 19.38 |
|  | Onit74 | Iran | Azarbaijan West | Sar dasht | 30.33 | 50.22 |
|  | Onit119 | Iran | Azarbaijan West | Arasbaran | 38.91 | 46.87 |
| Ebru | Ebru719 | Spain | - | Navacerrada | 40.73 | -4.00 |
|  | Ebru1132 | Spain | - | Nuevo Baztan | 40.37 | -3.24 |
|  | Ebru1927 | Hungary | - | Márkó | 47.12 | 17.81 |
|  | Ebru989 | Hungary | - | Godollo | 47.60 | 19.38 |
|  | Ebru664 | Iran | Kordestan | Marivan | 35.52 | 46.17 |
|  | Ebru2453 | Iran | Azerbaijan East | East Azerbaijan | 38.08 | 46.28 |
| Eann | Eann76 | Spain | - | Fresnedoso & Sorihuela | 40.43 | -5.70 |
|  | Eann69 | Spain | Ávila | La Cañada | 40.60 | -4.48 |
|  | Eann86 | Hungary | - | Várpolata | 47.20 | 18.15 |
|  | Eann87 | Hungary | - | Várpolata | 47.20 | 18.15 |
|  | Eann38 | Iran | Kermanshah | Gahvareh | 34.35 | 46.42 |
|  | Eann89 | Iran | Kordestan | Bane | 35.99 | 45.90 |
| Taur | Taur409 | Spain | - | El Escorial | 40.58 | -4.13 |
|  | Taur1384 | Portugal | - | Mirandela | 41.49 | -7.18 |
|  | Taur130 | Hungary | - | Jaszbereny | 47.50 | 19.92 |
|  | Taur246 | Hungary | - | Jaszbereny | 47.50 | 19.92 |
|  | Taur28 | Iran | Kordestan | Marivan | 35.52 | 46.17 |
|  | Taur263 | Iran | Lorestan | Ghelaie | 33.78 | 47.93 |
| Sumb | Inq2276 | Spain | Albacete | Carboneras | 37.00 | -1.88 |
|  | Inq2277 | Spain | Albacete | Carboneras | 37.00 | -1.88 |
|  | Sumb541 | Hungary | - | Orfű | 46.13 | 18.15 |
|  | Sumb488 | Hungary | - | Pécsvárad | 46.07 | 18.23 |
|  | Sumb623 | Iran | Azerbaijan West | Piranshahr | 36.70 | 45.14 |
|  | Inq2244 | Iran | Lorestan | Ghelaie | 33.78 | 47.93 |

**Table S3.**
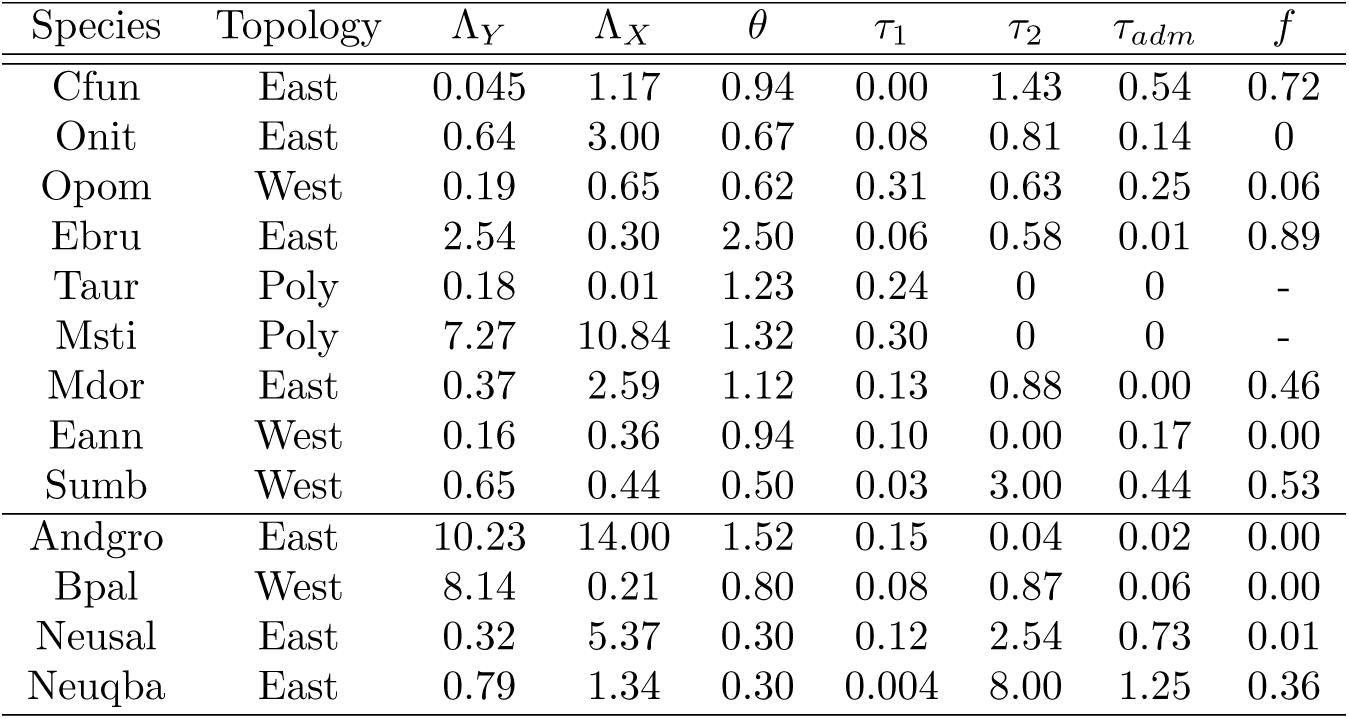
Uncalibrated parameter estimates for best supported models for each species. *θ* is the scaled mutation rate relating to the *N_e_* of the older population (i.e., the Eastern population for Out of the East and polytomous species and the Western population for Out of the West species). *τ* parameters are in coalescent units and are specified as intervals rather than from the present (i.e., the sum of *τ_adm_*, *τ*_1_ & *τ*_2_ gives the oldest split time in coalescent units. Λ parameters are scalars for N_e_ estimates for the remaining populations (Y = the central population, X = the Western population for Out of the East/polytomous species and the Eastern population for Out of the West species). See Table 1 for calibrated parameter estimates and species abbreviations.

| Species | Topology | $\Lambda_Y$ | $\Lambda_X$ | $\theta$ | $\tau_1$ | $\tau_2$ | $\tau_{adm}$ | $f$ |
| --- | --- | --- | --- | --- | --- | --- | --- | --- |
| Cfun | East | 0.045 | 1.17 | 0.94 | 0.00 | 1.43 | 0.54 | 0.72 |
| Onit | East | 0.64 | 3.00 | 0.67 | 0.08 | 0.81 | 0.14 | 0 |
| Opom | West | 0.19 | 0.65 | 0.62 | 0.31 | 0.63 | 0.25 | 0.06 |
| Ebru | East | 2.54 | 0.30 | 2.50 | 0.06 | 0.58 | 0.01 | 0.89 |
| Taur | Poly | 0.18 | 0.01 | 1.23 | 0.24 | 0 | 0 | - |
| Msti | Poly | 7.27 | 10.84 | 1.32 | 0.30 | 0 | 0 | - |
| Mdor | East | 0.37 | 2.59 | 1.12 | 0.13 | 0.88 | 0.00 | 0.46 |
| Eann | West | 0.16 | 0.36 | 0.94 | 0.10 | 0.00 | 0.17 | 0.00 |
| Sumb | West | 0.65 | 0.44 | 0.50 | 0.03 | 3.00 | 0.44 | 0.53 |
| Andgro | East | 10.23 | 14.00 | 1.52 | 0.15 | 0.04 | 0.02 | 0.00 |
| Bpal | West | 8.14 | 0.21 | 0.80 | 0.08 | 0.87 | 0.06 | 0.00 |
| Neusal | East | 0.32 | 5.37 | 0.30 | 0.12 | 2.54 | 0.73 | 0.01 |
| Neuqba | East | 0.79 | 1.34 | 0.30 | 0.004 | 8.00 | 1.25 | 0.36 |

**Table S4.** Identifying clusters of co-diverged species The best supported model without a significant decrease in *ΔlnL* (indicated in bold) contains two clusters of co-diverged species for *T*_2_. *ΔlnL* = *ΔlnL* between a model where all species have their own divergence time and one where the specified species co-diverge (see Table 1 for species abbreviations).

| No. of clusters | Species | $\Delta \ln L$ |
| --- | --- | --- |
| 1 | *Mdor*, *Bpal* | 0.09 |
| 1 | *Taur*, *Mdor*, *Bpal* | 1.47 |
| **2** | ***Taur*, *Mdor*, *Bpal* +** | **3.15** |
|  | ***Andgro*, *Msti*** |  |
| 3 | *Mdor*, *Bpal* + | 4.22 |
|  | *Andgro*, *Msti* + |  |
|  | *Cfun*, *Ebru* |  |

### Supp. Info 3 Assessing the power of the inference framework

To assess the power of our composite likelihood framework to distinguish between models we conducted a simple simulation study: we picked one of the 48 possible models at random and chose a random set of model parameters (by picking parameter values uniformly from the range of observed estimates). Applying our model selection to 100 such datasets, each simulated for a random point in model and parameter space, showed that model selection based on lnCL can identify the correct topology and admixture direction in 100 % of cases and the correct relationship of ancestral and current N_e_ parameters in 75 % of cases.

## References

1 Ricklefs RE. Intrinsic dynamics of the regional community. Ecology Letters. 2015;18(6):497–503. Available from: http://dx.doi.org/10.1111/ele.12431.

2 Jousselin E, Rasplus JY, Kjellberg F. Convergence and coevolution in a mutualism: evidence from a molecular phylogeny of Ficus. Evolution. 2003;57(6):1255–1269. Available from: http://dx.doi.Org/10.1111/j.0014-3820.2003.tb00334.x.

3 Mueller UG, Rehner SA, Schultz TR. The evolution of agriculture in ants. Science. 1998;281(5385):2034–2038. Available from: http://science.sciencemag.org/content/281/5385/2034.

4 Stone GN, Schönrogge K, Atkinson RJ, Bellido D, Pujade-Villar J. The population biology of oak gall waps (Hymenoptera: Cynipidae). Annual Review of Entomology. 2002;47(1):633–668.

5 Leppänen SA, Altenhofer E, Liston AD, Nyman T. Ecological versus phylogenetic determinants of trophic associations in a plant-leafminer-parasitoid food web. Evolution. 2013;67(5):1493–1502. Available from: http://dx.doi.org/10.1111/evo.12028.

6 Bailey R, Schönrogge K, Cook JM, Melika G, Csoka G, Thúroczy C, et al. Host niches and defensive extended phenotypes structure parasitoid wasp communities. PLoS Biology. 2009;7(8):e1000179.

7 Askew RR. The diversity of insect communities in leaf mines and plant galls. The Journal of Animal Ecology. 1980;49:145–152.

8 Stireman JO, Singer MS. Determinants of parasitoid-host associations: insights from a natural tachinid-lepidopteran community. Ecology. 2003;84(2):296–310. Available from: http://dx.doi.org/10.1890/0012-9658(2003)084[0296:D0PHAI]2.0.C0;2.

9 Hubbell SP. The unified neutral theory of biodiversity and biogeography. Princeton Monographs in Population Biology. Princeton, New Jersey, USA: Princeton University Press; 2001.

10 Rosindell J, Hubbell SP, Etienne RS. The unified neutral theory of biodiversity and biogeography at age ten. Trends in Ecology & Evolution. 2011;26(7):340 - 348. Available from: http://www.sciencedirect.com/science/article/pii/S0169534711000942.

11 Lohse K, Sharanowski B, Stone GN. Quantifying the population history of the oak gall parasitoid *Cecidostiba fungosa*. Evolution. 2010;58(4):439–442.

12 Stone GN, Challis RJ, Atkinson RJ, Csóka G, Hayward A, Melika G, et al. The phylogeographical clade trade: tracing the impact of human-mediated dispersal on the colonization of northern Europe by the oak gallwasp *Andricus kollari*. Molecular Ecology. 2007;16:2768–2781.

13 Hayward A, Stone GN. Comparative phylogeography across two trophic levels: the oak gall wasp *Andricus kollari* and its chalcid parasitoid *Megastigmus stigmatizans*. Molecular Ecology. 2006;15(2):479–489.

14 Nicholls JA, Preuss S, Hayward A, Melika G, Csóka G, Nieves-Aldrey JL, et al. Concordant phylogeography and cryptic speciation in two Western Palaearctic oak gall parasitoid species complexes. Molecular Ecology. 2010;19:592–609.

15 Stone GN, Hernandez-Lopez A, Nicholls JA, di Pierro E, Pujade-Villar J, Melika G, et al. Extreme host plant conservatism during at least 20 million years of host plant pursuit by oak gallwasps. Evolution. 2009;63(4):854–869.

16 Stone GN, Lohse K, Nicholls JA, Fuentes-Utrilla P, Sinclair F, Schönrogge K, et al. Reconstructing community assembly in time and space reveals enemy escape in a western palaearctic insect community. Current Biology. 2012;22(6):531–537.

17 Prior KM, Hellmann JJ. Does enemy loss cause release? A biogeographical comparison of parasitoid effects on an introduced insect. Ecology. 2013;94(5):1015–1024. Available from: http://dx.doi.org/10.1890/12-1710..

18 Lohse K, Chmelik M, Martin SH, Barton NH. Efficient strategies for calculating blockwise likelihoods under the coalescent. Genetics. 2016;202(2):775–786. Available from: http://www.genetics.org/content/202/2Z775.

19 Lohse K, Harrison RJ, Barton NH. A general method for calculating likelihoods under the coalescent process. Genetics. 2011;58(189):977–987.

20 Romiguier J, Gayral P, Ballenghien M, Bernard A, Cahais V, Chenuil A, et al. Comparative population genomics in animals uncovers the determinants of genetic diversity. Nature. 2014;515:261–263.

21 Leffler EM, Bullaughey K, Matute DR, Meyer WK, Ségurel L, Venkat A, et al. Revisiting an old riddle: what determines genetic diversity levels within species? PLOS Biology. 2012 09;10(9):1–9. Available from: https://doi.org/10.1371/journal.pbio.1001388.

22 Keightley PD, Trivedi U, Thomson M, Oliver F, Kumar S, Blaxter ML. Analysis of the genome sequences of three *Drosophila melanogaster* spontaneous mutation accumulation lines. Genome Research. 2009;19(7):1195–1201.

23 Stone GN, van der Ham RWJM, Brewer JG. Fossil oak galls preserve ancient multitrophic interactions. Proceedings of the Royal Society B: Biological Sciences. 2008;275(1648):2213–2219.

24 Lohse K, Barton N N H Melika, Stone GN. Likelihood based inference of of an guild. Molecular Ecology. 2012;49(3):832–842.

25 Ricklefs RE. Disintegration of the Ecological Community. The American Naturalist. 2008;172(6):741–750. PMID: 18954264.

26 Askew RR. On the biology of the inhabitants of oak galls of Cynipidae (Hymenoptera) in Britain. Transactions of the Society for British Entomology. 1961;14:237–258.

27 Corbett-Detig RB, Hartl DL, Sackton TB. Natural Selection Constrains Neutral Diversity across A Wide Range of Species. PLOS Biology. 2015 04;13(4):1–25. Available from: https://doi.org/10.1371/journal.pbio.1002112.

28 Koch MA, Kiefer C, Ehrich D, Vogel J, Brochmann C, Mummenhoff K. Three times out of Asia Minor: the phylogeography of Arabis alpina L. (Brassicaceae). Molecular Ecology. 2006;15(3):825–839. Available from: http://dx.doi.org/10.1111/j.1365-294X.2005.02848.x.

29 Duvaux L, Belkhir K, Boulesteix M, Boursot P. Isolation and gene flow: inferring the speciation history of European house mice. Molecular Ecology. 2011;20(24):5248–5264. Available from: http://dx.doi.org/10.1111/j.1365-294X.2011.05343.x.

30 Chapman JW, Reynolds DR, Smith AD, Smith ET, Woiwod IP. An aerial netting study of insects migrating at high altitude over England. Bulletin of Entomological Research. 2004;94(2):123–136.

31 Hickerson MJ, Stahl EA, Lessios HA, Crandall K. Test for simultaneous divergence using approximate Bayesian computation. Evolution. 2006;60(12):2435–2453.

32 Hearn J, Stone GN, Bunnefeld L, Nicholls JA, Barton NH, Lohse K. Likelihood-based inference of population history from low-coverage de novo genome assemblies. Molecular Ecology. 2014;23(1):198–211. Available from: http://dx.doi.org/10.1111/mec.12578.

33 Yu Y, Dong J, Liu KJ, Nakhleh L. Maximum likelihood inference of reticulate evolutionary histories. Proceedings of the National Academy of Sciences. 2014;111(46):16448–16453. Available from: http://www.pnas.org/content/111/46/16448.abstract.

34 Marko PB, Hart MW. The complex analytical landscape of gene flow inference. Trends in Ecology & Evolution. 2011;26(9):448 - 456.

35 Hung CM, Drovetski SV, Zink RM. The roles of ecology, behaviour and effective population size in the evolution of a community. Molecular Ecology. 2017;26(14):3775–3784. Available from: http://dx.doi.org/10.1111/mec.14152.

36 Sutton TL, Riegler M, Cook JM. One step ahead: a parasitoid disperses farther and forms a wider geographic population than its fig wasp host. Molecular Ecology. 2016;25(4):882–894. Available from: http://dx.doi.org/10.1111/mec.13445.

37 Espíndola A, Carstens BC, Alvarez N. Comparative phylogeography of mutualists and the effect of the host on the genetic structure of its partners. Biological Journal of the Linnean Society. 2014;113(4):1021–1035. Available from: +http://dx.doi.org/10.1111/bij.12393.

38 Beeravolu Reddy C, Hickerson MJ, Frantz LAF, Lohse K. Blockwise site frequency spectra for inferring complex population histories and recombination. bioRxiv. 2017; Available from: https://www.biorxiv.org/content/early/2017/09/25/077958.

39 Xue AT, Hickerson MJ. The aggregate site frequency spectrum for comparative population genomic inference. Molecular Ecology. 2015;24(24):6223–6240. Available from: http://dx.doi.org/10.1111/mec.13447.

40 Papadopoulou A, Knowles LL. Toward a paradigm shift in comparative phylogeography driven by trait-based hypotheses. Proceedings of the National Academy of Sciences. 2016;113(29):8018–8024. Available from: http://www.pnas.org/content/113/29/8018.abstract.

41 Satler JD, Carstens BC. Do ecological communities disperse across biogeographic barriers as a unit? Molecular Ecology. 2017;26(13):3533–3545. Available from: http://dx.doi.org/10.1111/mec.14137.

42 Oswald JA, Overcast I, Mauck WM, Andersen MJ, Smith BT. Isolation with asymmetric gene flow during the nonsynchronous divergence of dry forest birds. Molecular Ecology. 2017;26(5):1386–1400. Available from: http://dx.doi.org/10.1111/mec.14013.

43 Rougemont Q, Gagnaire PA, Perrier C, Genthon C, Besnard AL, Launey S, et al. Inferring the demographic history underlying parallel genomic divergence among pairs of parasitic and nonparasitic lamprey ecotypes. Molecular Ecology. 2017;26(1):142–162. Available from: http://dx.doi.org/10.1111/mec.13664.

44 Bunnefeld L, Frantz LAF, Lohse K. Inferring bottlenecks from genome-wide samples of short sequence blocks. Genetics. 2015;201(3):1157–1169. Available from: http://www.genetics.org/content/201/3Z1157.

## Methods references

[M1] Bolger AM, Lohse M, Usadel B. Trimmomatic: A flexible trimmer for Illumina sequence data. Bioinformatics. 2014;30(15):2114–2120.

[M2] Martin M. Cutadapt removes adapter sequences from high-throughput sequencing reads. EMBnet journal. 2011;17(1):pp-10.

[M3] Andrews S. FastQC: A quality control application for high throughput sequence data. Available at: http://www.bioinformatics.babraham.ac.uk/projects/fastqc; 2010.

[M4] Bankevich A, Nurk S, Antipov D, Gurevich Aa, Dvorkin M, Kulikov AS, et al. SPAdes: a new genome assembly algorithm and its applications to single-cell sequencing. Journal of computational biology: a journal of computational molecular cell biology. 2012;19(5):455–77. Available from: http://www.pubmedcentral.nih.gov/articlerender.fcgi?artid=3342519{}tool=pmcentrez{}rendertype=abstract.

[M5] Zimin AV, Marçais G, Puiu D, Roberts M, Salzberg SL, Yorke Ja. The MaSuRCA genome assembler. Bioinformatics (Oxford, England). 2013;29(21):2669–77. Available from: http://www.pubmedcentral.nih.gov/articlerender.fcgi?artid=3799473{&}tool=pmcentrez{}rendertype=abstract.

[M6] Price AL, Jones NC, Pevzner PA. De novo identification of repeat families in large genomes. Bioinformatics. 2005;21(suppl 1):i351–i358. Available from: http://bioinformatics.oxfordjournals.org/content/21/suppl{_}1/i351.abstract.

[M7] Tarailo-Graovac M, Chen N. Using RepeatMasker to identify repetitive elements in genomic sequences. Current protocols in bioinformatics/editoral board, Andreas D Baxevanis [et al]. 2009;Chapter 4(March):Unit 4.10. Available from: http://www.ncbi.nlm.nih.gov/pubmed/19274634.

[M8] Altschul SF, Madden TL, Schaffer AA, Zhang J, Zhang Z, Miller W, et al. Gapped BLAST and PSI-BLAST: a new generation of protein database search programs. Nucleic Acids Research. 1997;25. Available from: http://dx.doi.org/10.1093/nar/25.17.3389.

[M9] Li H, Durbin R. Fast and accurate short read alignment with Burrows-Wheeler transform. Bioinformatics. 2009;25(14):1754–1760.

[M10] Auwera GAVD, Carneiro MO, Hartl C, Poplin R, Levy-Moonshine A, Jordan T, et al. From FastQ data to high confidence variant calls: the Genome Analysis Toolkit best practices pipeline. vol. 11; 2014.

[M11] Wolfram Research I. Mathematica, Version 11.1.1.0. Champaign, Illinois: Wolfram Research, Inc.; 2017.

[M12] Than C, Ruths D, Nakhleh L. PhyloNet: a software package for analyzing and reconstructing reticulate evolutionary relationships. BMC Bioinformatics. 2008;9(1):322. Available from: http://dx.doi.org/10.1186/1471-2105-9-322.

[M13] Costa RJ, Wilkinson-Herbots H. Inference of gene flow in the process of speciation: an efficient maximum-likelihood method for the isolation-with-initial-migration model. Genetics. 2017;205(4):1597–1618. Available from: http://www.genetics.org/content/205/4/1597.

[M14] Yang Z. A likelihood ratio test of speciation with gene flow using genomic data. Genome Biology and Evolution. 2010;2:200–211.

[M15] Pickrell JK, Pritchard JK. Inference of population splits and mixtures from genome-wide allele frequency data. PLOS Genetics. 2012 11;8(11):1–17. Available from: https://doi.org/10.1371/journal.pgen.1002967.

[M16] Kelleher J, Etheridge AM, McVean G. Efficient coalescent simulation and genealogical analysis for large sample sizes. PLOS Computational Biology. 2016 05;12(5):1–22. Available from: http://dx.doi.org/10.1371%2Fjournal.pcbi.1004842.

## Supplementary references

[S1] Nei M. Genetic distance between populations. The American Naturalist. 1972;106(949):283–292. Available from: https://doi.org/10.1086/282771.

[S2] Durand EY, Patterson N, Reich D, Slatkin M. Testing for ancient admixture between closely related populations. Molecular Biology and Evolution. 2011;28(8):2239–2252.

[S3] Askew RR, Melika G, Pujade-Villar J, Schönrogge K, Stone GN, Nieves-Aldrey JL. Catalogue of parasitoids and inquilines in cynipid oak galls in the West Palaearctic. Zootaxa. 2013;3643:1–133.

[S4] Orme D, Freckleton R, Thomas G, Petzoldt T, Fritz S, Isaac N, et al.. caper: comparative analyses of phylogenetics and evolution in R.; 2013. R package version 0.5.2.

[S5] Slater GSC, Birney E. Automated generation of heuristics for biological sequence comparison. BMC bioinformatics. 2005;6:31. Available from: http://www.pubmedcentral.nih.gov/articlerender.fcgi?artid=553969{}tool=pmcentrez{&}rendertype=abstract.

[S6] Quinlan AR, Hall IM. BEDTools: a flexible suite of utilities for comparing genomic features. Bioinformatics. 2010;26(6):841–842.

[S7] Abascal F, Zardoya R, Telford MJ. TranslatorX: multiple alignment of nucleotide sequences guided by amino acid translations. Nucleic acids research. 2010;38(suppl_2):W7–W13.

[S8] Thornton K. libsequence: A C++ class library for evolutionary genetic analysis. Bioinformatics. 2003;19(17):2325–2327.

[S9] Nordborg M, Charlesworth B, Charlesworth D. The effect of recombination on background selection. Genetical Research. 1996;67(2):159–174.

